# Efficacy of foramen magnum decompression with and without cranioplasty in a rat model of Chiari-like malformation

**DOI:** 10.1101/2024.04.23.590717

**Authors:** Jae-Hwan Jung, Chang-Hyeon Cho, Woo-Suk Kim, Hun-Young Yoon

**Affiliations:** Department of Veterinary Surgery, College of Veterinary Medicine, Konkuk University, 120 Neungdong-ro, Gwangjin-gu, Seoul 05029, Republic of Korea; Department of Veterinary Radiology, College of Veterinary Medicine, Konkuk University, 120 Neungdong-ro, Gwangjin-gu, Seoul 05029, Republic of Korea; Department of Anatomy, College of Veterinary Medicine, and Veterinary Science Research Institute, Konkuk University, 120 Neungdong-ro, Gwangjin-gu, Seoul 05029, Republic of Korea; KU Center for Animal Blood Medical Science, Konkuk University, 120 Neungdong-ro, Gwangjin-gu, Seoul 05029, Republic of Korea

**Author notes:** Corresponding author: (H-Y Y).

## Abstract

In veterinary medicine, canine Chiari-like malformation (CLM) disease is surgically managed through foramen magnum decompression (FMD) with cranioplasty. This study aimed to assess the efficacy of cranioplasty surgery by establishing a rat CLM model and then applying FMD with and without cranioplasty and comparing the outcomes. Twenty-four 8-week-old male Sprague‒Dawley rats underwent surgery to induce CLM by reducing the caudal cranial fossa volume, mimicking cerebellum herniation. The rats were randomly and equally assigned to three groups: a control group (induced CLM), an FO group (induced CLM rats undergoing FMD only), and a CR group (induced CLM rats undergoing FMD with cranioplasty). At 11 weeks of age, the FO and CR groups underwent FMD surgery. Four weeks later, magnetic resonance imaging (MRI) was used to measure the cisterna magna volume to assess surgical outcomes. Post-surgery MRI revealed that the mean cisterna magna volume was 23.82 ± 1.70, 34.88 ± 4.39, and 29.48 ± 2.20 mm^3^ in the control, FO, and CR groups, respectively. There was a significant increase in the cisterna magna volume in the FO and CR groups compared to that in the control group (p < 0.05), with the FO group showing a significantly greater increase than the CR group (p < 0.05). These findings suggest that FMD surgery alone is more effective at restoring the cisterna magna volume than FMD surgery with cranioplasty. FMD surgery alone resulted in a greater increase in cisterna magna volume than FMD with cranioplasty in our rat CLM model, suggesting that FMD alone may be more effective to treat canine CLM. These findings challenge the typical approach of combining FMD with cranioplasty in treating canine CLM disease and underscore the need for further investigation into optimizing surgical techniques for CLM.

## Introduction

Chiari-like malformation (CLM) predominantly affects Cavalier King Charles Spaniels and is known as Chiari Type 1 malformation (CM1) in humans [1–4]. This condition is characterized by a morphologically malformed skull and craniocervical junction, resulting in a relatively small caudal cranial fossa (CCF) and caudal displacement of the cerebellum into the foramen magnum due to overcrowding of the CCF [1–4]. Herniation of the cerebellum due to skull insufficiency causes deformation of the cerebellum, prompted by insufficient skull capacity, leading to deformation of the craniocervical junction, including loss of the cisterna magna. This deformation may subsequently cause syringomyelia by impeding the normal flow of cerebrospinal fluid (CSF), along with gradual constriction [5,6]. The clinical manifestations of CLM include pain, phantom scratching, head tilt, head tremor, loss of balance, tetraparesis, muscle atrophy, and scoliosis [6]. These symptoms are believed to stem from altered pressure dynamics between the intracranial and spinal compartments due to CSF pathway obstructions [2,6].

Although the features of the cisterna magna are less understood in veterinary medicine, recent studies in human medicine have suggested that headaches, a primary clinical symptom of CM1, may be linked to an anatomical feature known as a limited cisterna magna [7]. Located at the craniospinal junction, the cisterna magna lies inferior to the cerebellum and medulla and dorsal to the upper cervical spinal cord [8]. Like other cranial cisterns, the cisterna magna maintains a wide CSF compartment, regulating rapid pressure changes within the cranial and spinal compartments during physiological and pathological events [8]. However, in CM1, decreased posterior cranial fossa volume leads to herniation of the cerebellar tonsils, which invade the cisterna magna space, disrupting CSF flow and leading to clinical symptoms [7].

Treatment options include medical and surgical treatment through foramen magnum decompression (FMD). However, the underlying pathophysiology of CLM remains poorly understood; thus, no standardized treatment approach exists [2]. While medical treatment can initially alleviate pain, its effectiveness may decrease, even with increased dosages and additional medications [2]. Therefore, surgical intervention to address structural abnormalities, the root cause of CLM, is recommended [9]. In addition to FMD, commonly used to treat human CM1, veterinary medicine often employs cranioplasty using titanium mesh/polymethyl-methacrylate (PMMA) plates [10]. However, cranioplasty only provides temporary relief of symptoms [3]. Moreover, the short-term surgical success rate for FMD and FMD with cranioplasty in patients with CLM is approximately 80%, showing little difference between the two methods [9,11]. Determination of the prognosis is, therefore, challenging due to the lack of long-term follow-up data, with clinical symptoms often persisting or recurring postoperatively [9,11].

Given these considerations, we questioned whether combining FMD with cranioplasty, which aims to cover the bony defect created by FMD, is more effective than FMD alone. Therefore, this study aimed to create a rat CLM model and compare the efficacy of FMD with and without cranioplasty by assessing changes in the volume of the cisterna magna following surgery.

## Materials and methods

### Animals

Thirty-two male Sprague‒Dawley (SD) rats, weighing 210–230 g and aged 7 weeks, were acquired (DBL Co., Ltd., Chungbuk, Republic of Korea) and acclimatized to their new environment for 1 week. The rats were housed in individually ventilated cages at a temperature of 22 ± 2°C and 55 ± 5% relative humidity under a 12-h day/12-h night cycle with unrestricted access to water and food. This study was approved by the Institutional Animal Care and Use Committee of Konkuk University (Approval number KU23177).

### Creating a rat CLM model before the main experiment

To create a rat CLM model, twenty-four 8-week-old male SD rats weighing 240–270 g were placed in a small induction chamber and anesthetized with 4% isoflurane in oxygen for 2 min. Following induction, the rats were transferred to a surgical table and maintained on 1–2% isoflurane through a nose hose [12]. The surgical area was sheared and sterilized using povidone and alcohol.

A skin incision of approximately 2 cm from the bregma to the atlas was made with a No. 15 blade. Upon revealing the periosteum covering the calvarium, a sagittal midline periosteal incision was made, and the periosteum was then elevated laterally with a periosteal elevator. The foramen magnum was exposed along the interparietal and occipital bones by separating the muscles attached to the occipital bone. From the diametric endpoints of the foramen magnum, a portion of the occipital and interparietal bones was vertically resected using a high-speed dental handpiece equipped with a 0.6 mm diameter carbide bur (FG 1/2, MANI Inc., Japan), extending 3 mm rostral from the external occipital protuberance and 2 mm caudal from the lambda (Fig 1A). The burring speed did not exceed 1,000 rpm, and constant irrigation with sterile normal saline was applied to prevent thermal damage [12]. Afterward, the removed bone fragment was placed in an NaCl (0.9%) solution.

**Fig 1.**
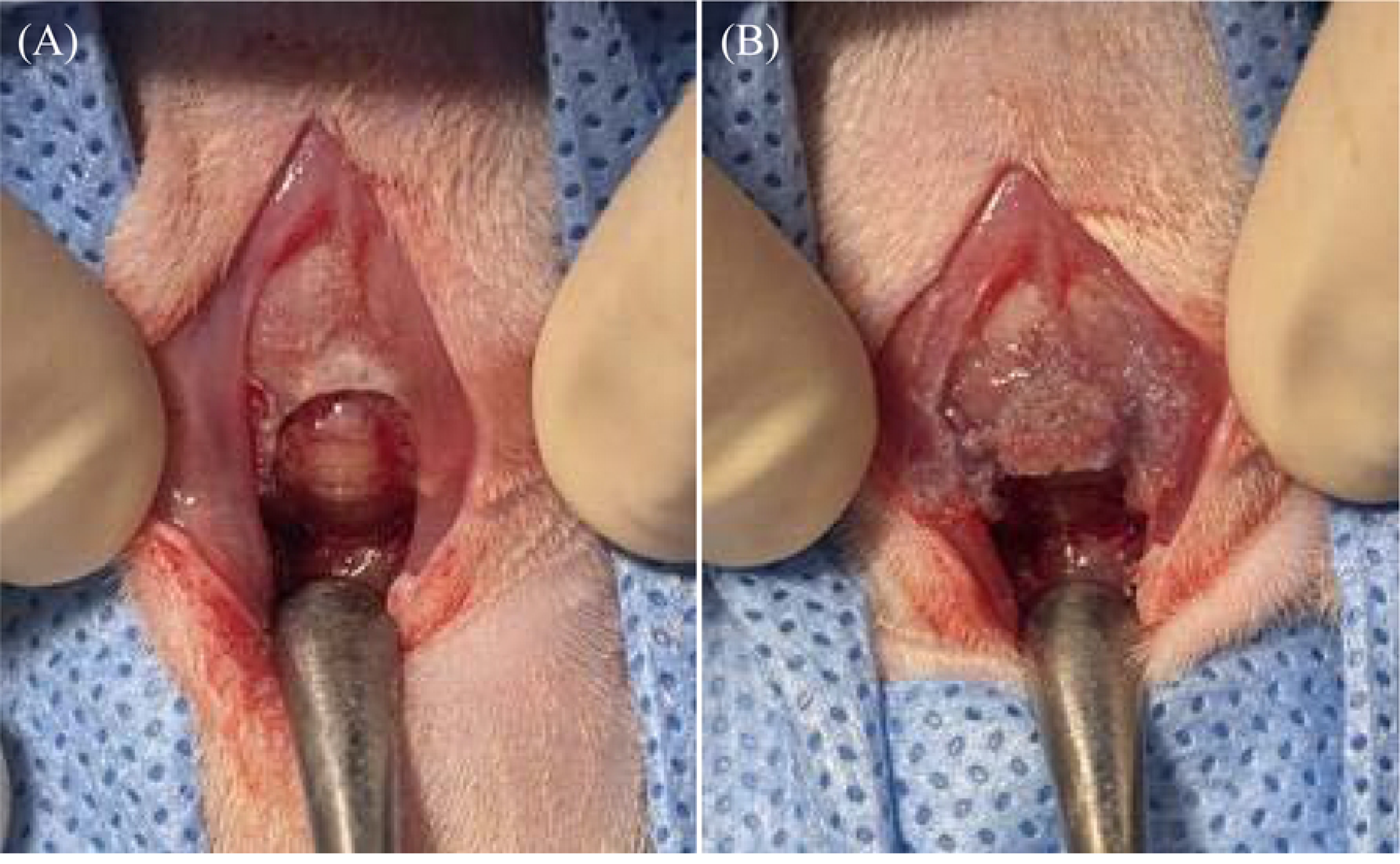
Creation of a rat Chiari-like malformation (CLM) model in an 8-week-old male rat (A) A carbide bur (FG 1/2, MANI Inc., Japan) was used to partially resect the occipital and interparietal bones. The resection was made vertically from the diametric endpoints of the foramen magnum, 3 mm rostral to the external occipital protuberance, and 2 mm caudal to the lambda. (B) A small hole, 0.6 mm in diameter, is drilled, and the 3 mm protruding part of the resected occipital bone fragment is positioned to overlap with the parietal and interparietal bone, constituting the skull.

The brain surface was regularly moistened with an NaCl solution. A small hole, 0.6 mm in diameter, was drilled 1 mm from the front border of the resected occipital bone fragment using a 0.6 mm diameter carbide bur (FG 1/2, MANI Inc., Japan). The 3-mm protruding part of the resected occipital bone was overlaid on the parietal and interparietal bones comprising the skull and secured using Vetbond™® (3M Company) (Fig 1B). The periosteum was sutured using simple interrupted stitches with 5-0 absorbable monofilament sutures (Monocryl®, Ethicon). Skin closure was performed with 4-0 nonabsorbable monofilament sutures (Dafilon®, B. Braun, Spain).

### MRI examination of the rat CLM model

Magnetic resonance imaging (MRI) scans were obtained 2 weeks post-surgery to assess the induction of CLM in the rats. The criteria for CLM formation were based on the Chiari Malformation grading guidelines of the British Veterinary Association to classify the severity of structural abnormalities in dogs [13]. MRI confirmed cerebellar herniation through the foramen magnum in all subjects, resulting in a diagnosis with CLM grade 2. Consequently, 24 rat CLM models were established (Fig 2).

**Fig 2.**
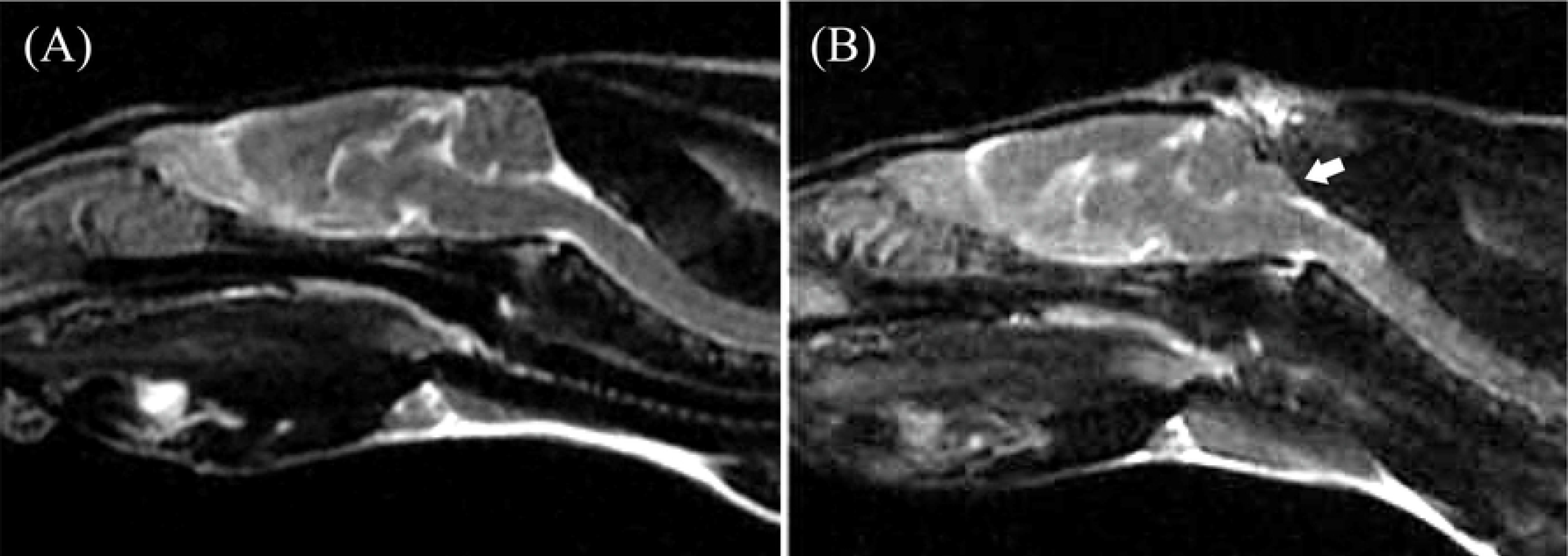
T2-weighted sagittal magnetic resonance imaging (MRI) of the brain of a 10-week-old male rat (A) Normal brain; (B) rat Chiari-like malformation model in which cerebellar herniation (white arrow) was induced into the foramen magnum by surgically reducing the caudal cranial fossa volume.

### Experimental groups

At 11 weeks, the 24 rat CLM models were randomly and equally divided into three groups: a control group with CLM (n = 8), a group with CLM undergoing FMD surgery (FO group, n = 8), and a group with CLM undergoing FMD with cranioplasty (CR group, n = 8). The rats weighed between 300 and 330 g.

All surgical procedures were performed under aseptic conditions. FMD surgery or FMD with cranioplasty surgery was performed on the 11-week-old rat CLM models, and an MRI was conducted 4 weeks later to evaluate the cisterna magna volume.

### Surgical procedure

#### FMD

The animals were placed in a small induction chamber and anesthetized with 4% isoflurane in oxygen for 2 min [12]. After induction, the rats were transferred to a surgical table and maintained on 1–2% isoflurane through a nose hose [12]. The rats were positioned in sternal recumbency, with the neck ventroflexed. The dorsal surface hair from the parietal bone to the third or fourth cervical vertebra, approximately the width of the first cervical vertebra, was shaved, and the area was sterilized.

A dorsal midline incision was made using a No. 15 blade, extending approximately 5 mm rostral from the external occipital protuberance to the middle of the second cervical vertebra. The superficial dorsal cervical musculature was separated from the median raphe to expose the underlying biventer cervicis muscle [14]. The paired biventer cervicis muscles were separated along the midline, revealing the rectus capitis dorsalis muscle. Sharp dissection and periosteal elevation were used to detach the caudal aspect of the rectus capitis dorsalis muscle from the cranial half of C2 and to separate the muscles in the midline. The cranial aspects of the rectus dorsalis muscle were sharply incised from the nuchal crest, exposing the atlas arch and the caudal portion of the occiput. Hemostasis was achieved with bipolar electrocautery.

To adequately expose the cerebellum, resection of part of the occipital bone was performed using an air-driven, high-speed dental handpiece equipped with a 0.6-mm diameter carbide bur (FG 1/2, MANI Inc., Japan), Kerrison rongeurs, and Lempert rongeurs (Fig 3). Subsequently, C1 laminectomy was performed using the same equipment.

**Fig 3.**
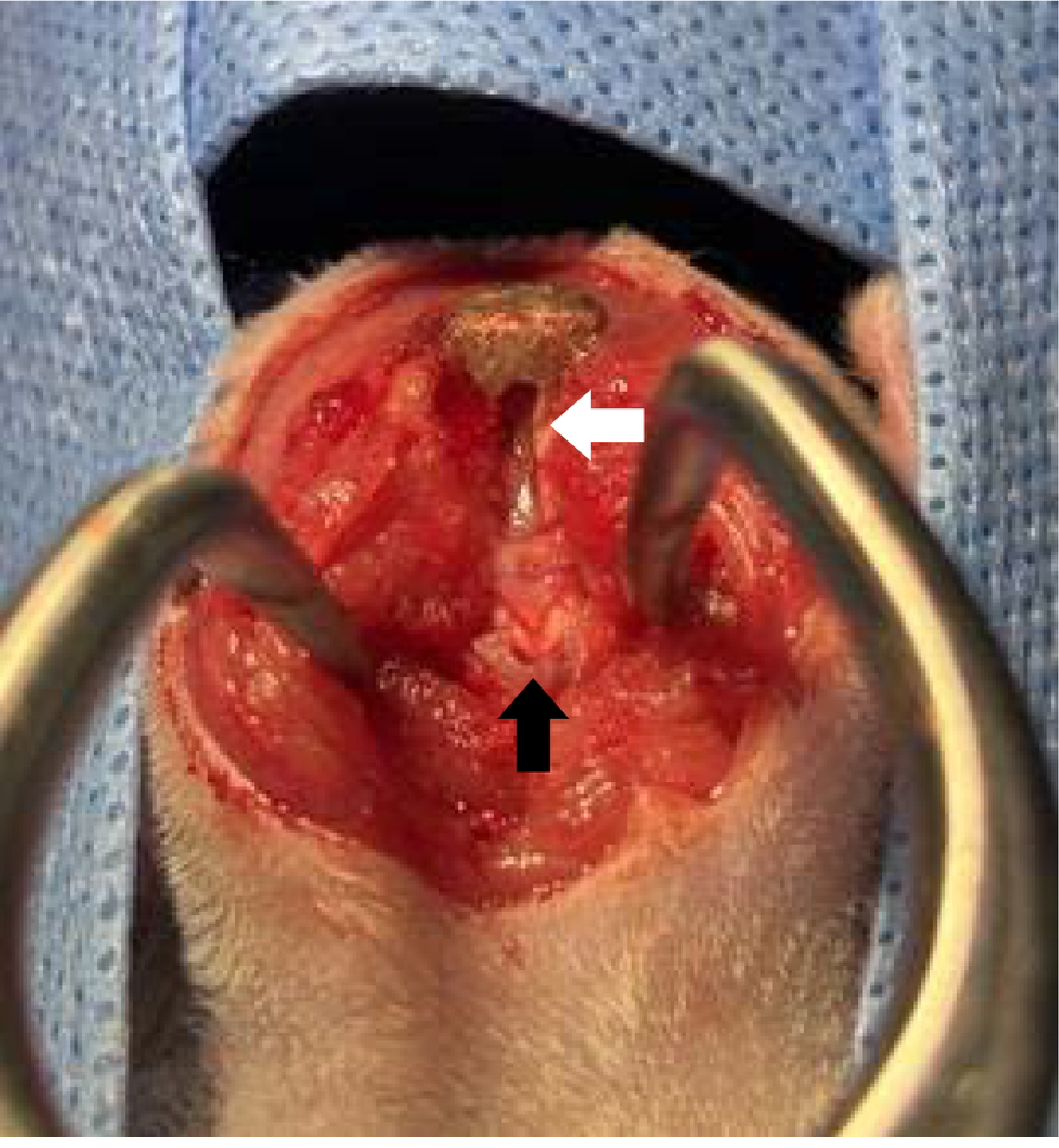
Surgical image of foramen magnum decompression (FMD) in a Chiari-like malformation (CLM) model in a male rat at 11 weeks of age Partial resection of the occipital bone was performed to expose the cerebellum (white arrow) adequately. This was followed by a C1 laminectomy, which removed approximately 75% of the length of the dorsal arch of the C1 (black arrow).

For the laminectomy, approximately 75% of the C1 dorsal arch length was removed (Fig 3). The muscle layers were sutured with 5-0 absorbable monofilament sutures (Monocryl®, Ethicon), and the skin was closed with 4-0 nonabsorbable monofilament sutures (Dafilon®, B. Braun, Spain). Postoperatively, cefotaxime (30 mg/kg, SC, once daily), enrofloxacin (5 mg/kg, SC, once daily), tramadol (10 mg/kg, SC, once daily), and meloxicam (2 mg/kg, SC, once daily) were administered for 2 weeks. The surgical site was disinfected daily with a 10% povidone-iodine solution for 2 weeks.

#### FMD and cranioplasty

In the CR group, the FMD procedure was conducted as described above, followed by cranioplasty. The rat’s head was returned to its normal resting angle after releasing it from the bent position. Titanium mesh (TM) was placed on the back of the skull, covering the site of the FMD surgery, and secured with Vetbond™ (3M Company) (Fig 4A).

**Fig 4.**
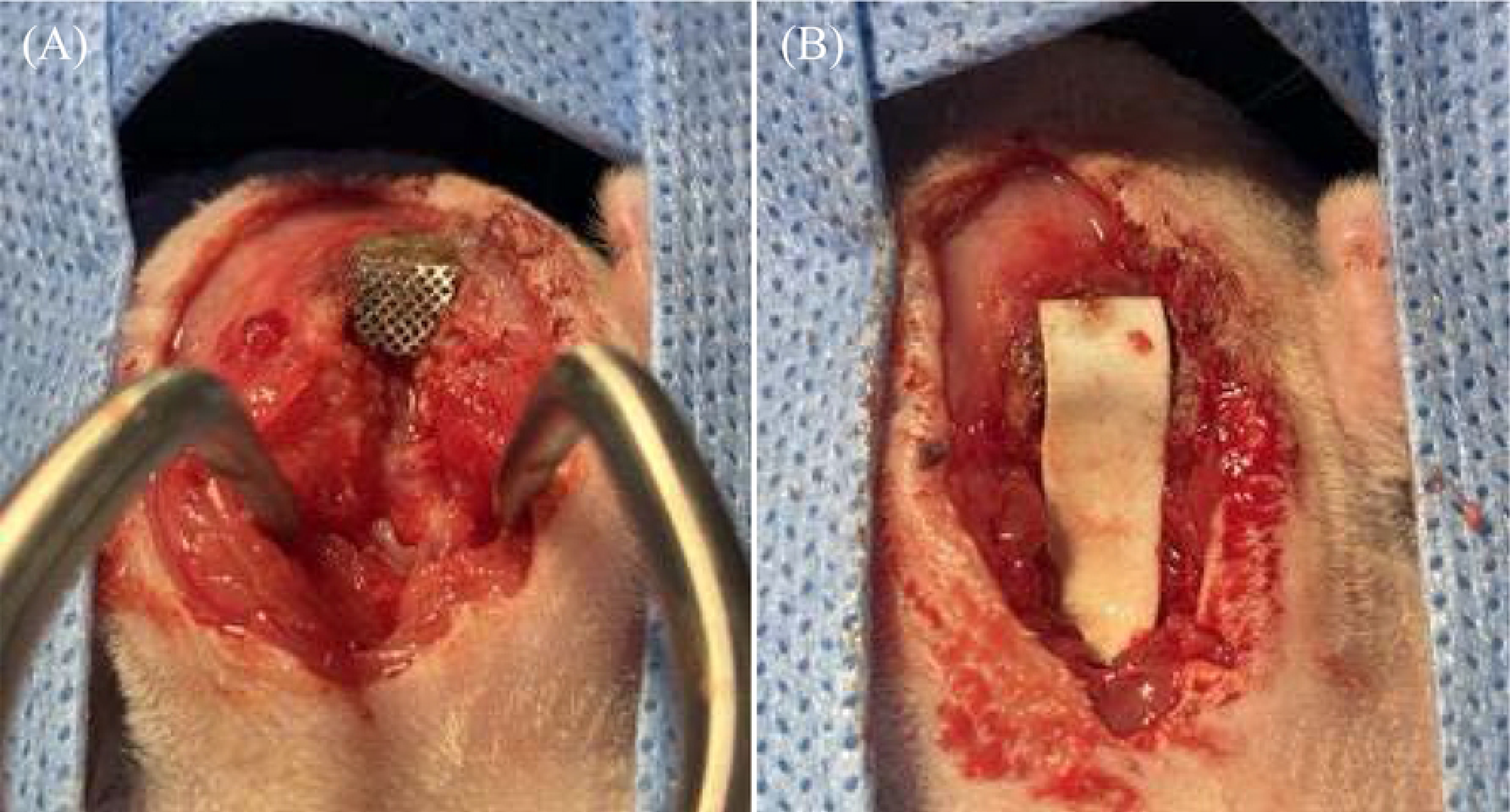
Cranioplasty on an 11-week-old male rat Chiari-like malformation (CLM) model (A) The titanium mesh (TM) is positioned over the caudal aspect of the skull, corresponding to the FMD surgical site, and secured in place. (B) Following the secure placement of the TM over the caudal aspect of the skull, reconstruction of the caudal occipital region is achieved by fabricating a skull plate using PMMA bone cement.

A skull plate was made using TM and PMMA bone cement. PMMA was used to mold a skull plate resembling a guitar pick, with the wide end facing toward the back of the head. A thin layer of PMMA was created on both sides of the skull plate, and a small amount of PMMA was placed beyond the edge of the TM and positioned on the caudal side of the skull plate, slightly beyond the dorsal defect at C1 (Fig 4B). Care was taken to bend the tail aspect of the headboard dorsally to avoid contact with the medulla oblongata or cranial cervical spinal cord. The muscle layers were sutured layer by layer using 5-0 absorbable monofilament sutures (Monocryl®, Ethicon), and skin closure was performed with 4-0 nonabsorbable monofilament sutures (Dafilon®, B. Braun, Spain).

Postoperative care included the administration of cefotaxime (30 mg/kg, SC, once daily), enrofloxacin (5 mg/kg, SC, once daily), tramadol (10 mg/kg, SC, once daily), and meloxicam (2 mg/kg, SC, once daily) for 2 weeks. The surgical site was disinfected daily with a 10% povidone-iodine solution for 2 weeks.

### MRI examination

The volume of the cisterna magna was measured using MRI at 4 weeks post-surgery for the FO and CR groups (Figs 5B, C). For the control group, MRI scans were performed at 15 weeks of age, aligning with the timing of the MRI scans in the surgical groups (Fig 5A).

**Fig 5.**
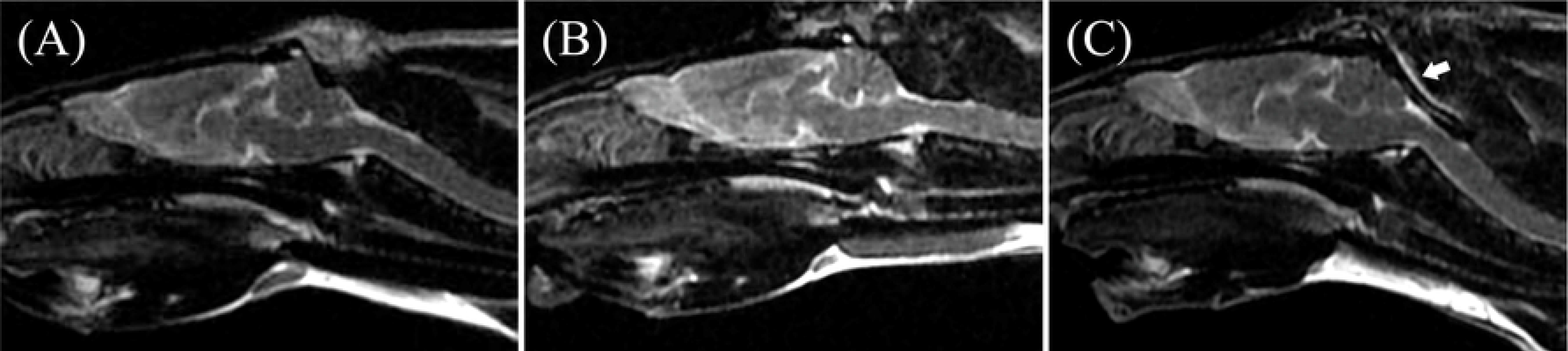
MRI images of a 15-week-old male rat Chiari-like malformation (CLM) model (A) A 15-week-old male rat CLM model in the control group. Cerebellar herniation into the foramen magnum was surgically induced by reducing the CCF volume. (B) A 15-week-old male rat CLM model in the FO group (FMD surgery only). (C) A 15-week-old male rat CLM model in the CR group (FMD with cranioplasty). The skull plate (white arrow) was molded with PMMA during cranioplasty surgery.

Cisterna magna volume measurements were conducted by the same board-certified radiologist. Each subject was assigned a study number, and details such as breed, age, sex, and weight were provided to the measurer in a blinded manner to facilitate volumetric measurements, thus minimizing measurer bias. All images were captured using a 1.5-T MRI (Signa Hdxt, GE) with a knee coil (eight channels). MRI sequences included a two-dimensional T2-weighted sagittal plane, a three-dimensional (3D) T2W CUBE transverse plane, and a 3D T1W FSPGR transverse plane. Volume rendering image processing was performed using image analysis and scientific visualization software (3D Slicer 5.4.0, 2023.8.19), which supports manual and semiautomated image segmentation and 3D volume rendering functions, including manual creation of image masking to enhance the precision of image segmentation [15,16].

Images were processed as follows. First, the MR image was segmented by slice, and voxels representing the anatomical structures of the CCF and rostral medial fossa in each image were identified. The rostral limit of the cisterna magna was defined as the lower part of the cerebellum; the caudal limit, the cranial boundary of C1; the dorsal limit, the point of intersection between the occipital bone and the cerebellum; and the ventral limit, the dorsal surface of the medulla oblongata. These boundaries delineated the total cisterna magna. In instances of cerebellar herniation, the space occupied by the herniated cerebellum was excluded from the total cisterna magna space, and the remaining space was identified as the free area of the cisterna magna (hereinafter referred to as the cisterna magna) for comparison in this study [7] (Figs 6A, B, C).

**Fig 6.**
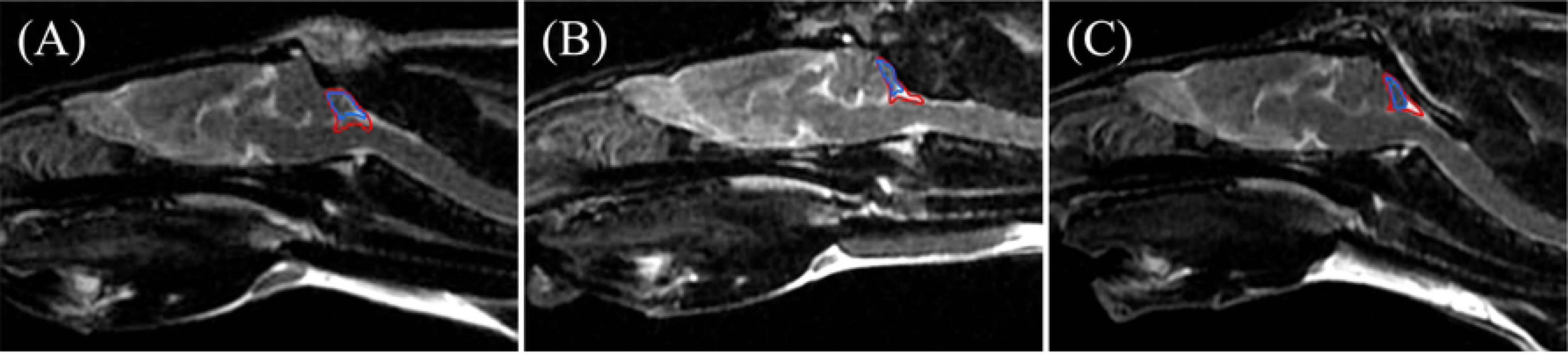
The free area of the cisterna magna in the groups The remaining space, after excluding the portion of the cisterna magna occupied by the herniated cerebellum from the total cisterna magna space, was defined as the free area of the cisterna magna (hereinafter referred to as the cisterna magna) to be compared in this experiment. The red line represents the boundary of the total cisterna magna space. The blue line represents the boundary of the space within the total cisterna magna occupied by the herniated cerebellum. (A) A 15-week-old male rat in the control group. (B) A 15-week-old male rat in the FO group. (C) A 15-week-old male rat in the CR group.

This methodology enhanced the accuracy of delineating the cisterna magna space. Subsequently, the segmented voxel values were masked to refine the delineation of anatomical structures [17]. The final mask incorporated data on all selected anatomical structures, and a 3D model was generated through volume rendering, integrating the original data with polygonal surface reconstruction algorithms [15, 17]. Measurement errors were minimized by quantifying the volume using image analysis and scientific visualization software (3D Slicer 5.4.0, 2023.8.19). The volume of the cisterna magna in the healthy, control, FO, and CR groups was measured twice, and the average volume was calculated for each group.

### Statistical analysis

Statistical analyses were performed using SPSS software (version 26.0, IBM Corp., Armonk, NY). The normality of the data distribution was assessed using the Shapiro‒Wilk test. Descriptive statistics, including the mean ± standard deviation and quartiles (Q1, Q2, Q3), are provided. Differences between the control, FO, and CR groups were analyzed using the Kruskal‒Wallis test, with post hoc testing performed using the Bonferroni correction. The significance level for all the tests was set at p < 0.05.

## Results

### MRI assessment

The volume of the cisterna magna was measured for all groups. Each volume measurement was taken twice, and the average value was calculated (Fig 7). The cisterna magna volume was 23.82 ± 1.70, 34.88 ± 4.39, and 29.48 ± 2.20 mm^3^ in the control, FO, and CR groups, respectively.

**Fig 7.**
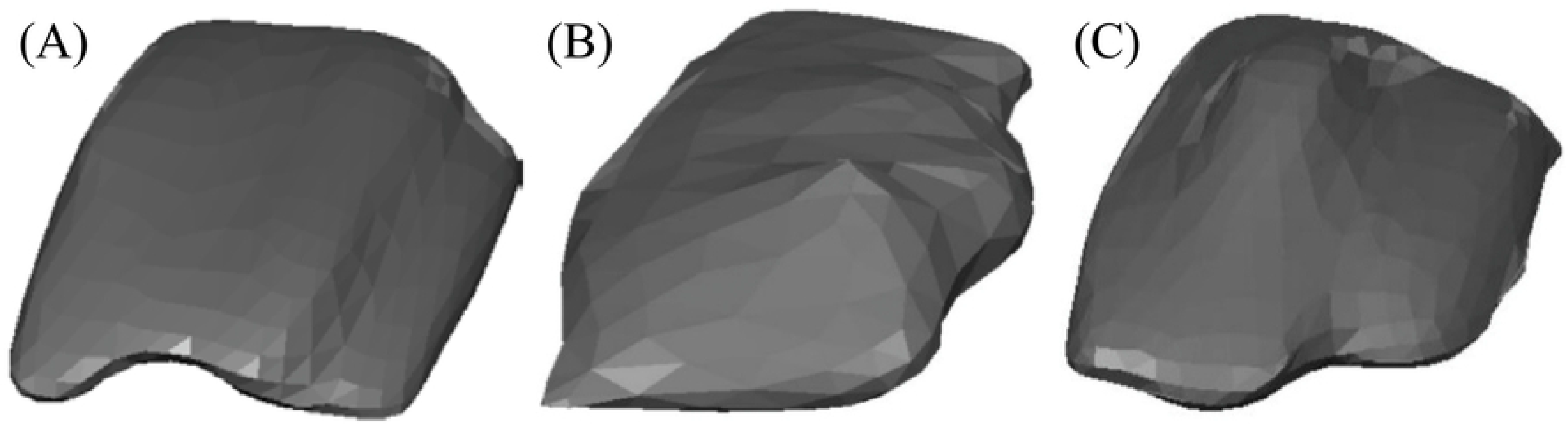
Dorsolateral view of the cisterna magna in 15-week-old male rat CLM models reconstructed using 3D volume rendering The cisterna magna volume in a 15-week-old male rat CLM model in the (A) control group, (B) FO group, and (C) CR group.

### Comparison of the cisterna magna volume among the groups

A comparison of cisterna magna volumes at 15 weeks among the control, FO, and CR groups is summarized in Table 1 (S1 Fig). The average cisterna magna volumes from the first and second measurements showed significant differences among the groups (p < 0.001). A post hoc analysis indicated that cisterna magna volumes were significantly greater in the FO group than in the control group (p < 0.001), the CR group than in the control group (p = 0.003), and the FO group than in the CR group (p = 0.005).

**Table 1.** Comparison of the cisterna magna volume between the control, FO, and CR groups.

|  | Control group | FO group | CR group |
| --- | --- | --- | --- |
| Cisterna magna (mm<sup>3</sup>) |  |  |  |
| Mean ± SD<sup>a</sup> | 23.82 ± 1.70 | 34.88 ± 4.39<sup>b</sup> | 29.48 ± 2.20<sup>b,c</sup> |
| (Median, Q1, Q3) | (23.44, 22.66, 25.48) | (35.42, 31.63, 37.96) | (29.60, 27.58, 30.18) |

<sup>a</sup>Post hoc Kruskal–Wallis test with Bonferroni correction.
<sup>b</sup>Significant difference compared to the control group ( $p < 0.05$ ).
<sup>c</sup>Significant difference compared to the FO group ( $p < 0.05$ ).
SD, standard deviation; FO group, foramen magnum decompression (FMD); CR group, FMD with cranioplasty.

## Discussion

This study compared FMD alone to FMD with cranioplasty in a rat CLM model, providing valuable insights into surgical treatments for CLM in veterinary medicine. The findings demonstrated that FMD alone and FMD with cranioplasty increased cisterna magna volume, suggesting that these methods are effective to treat CLM. Notably, FMD alone resulted in a larger increase in cisterna magna volume than that observed following FMD with cranioplasty, which is more commonly performed in clinical practice.

In humans, the cisterna magna is a large subarachnoid CSF space between the medulla oblongata and the lower portion of the cerebellum. CSF exits the fourth ventricle via the median and lateral apertures, flowing into the cisterna magna [18, 19]. In human CM1, herniation of the cerebellar tonsils into the cisterna magna at the foramen magnum displaces the subarachnoid space at the cervicomedullary junction and compromises this region [8]. Anatomical obstruction of the cisterna magna impedes CSF pulse pressure equalization and craniocaudal CSF flow between the intracranial and intraspinal compartments, elevating intracranial pressure and exerting pressure on adjacent neural structures [8]. This results in CM1 symptoms, including Valsalva-induced suboccipital headaches, upper and lower motor neuron signs, temperature sensory loss, spasticity, motor weakness, absence of the gag reflex, hoarseness, difficulties swallowing, and central sleep apnea [20].

Research on the cisterna magna has been limited in both veterinary and human medicine [18]. Only recently has the importance of the cisterna magna in the context of Chiari disease in humans been recognized, prompting further research, although there remains a lack of comprehensive knowledge [18]. A recent study highlighted that one primary objective of Chiari surgery is to restore free CSF flow through the functional restoration of the cisterna magna by performing posterior fossa decompression, addressing the pathology of obstructive CSF flow [8].

In human medicine, the anatomical boundaries of the cisterna magna are defined with the upper limit as the lower part of the cerebellum, the inferior border as the upper limit of C1, the anterior border as the inferior medullary velum and medulla, and the posterior border as the occipital bone [7]. Based on these findings, we established a new boundary for the cisterna magna in veterinary medicine, which, to the best of our knowledge, has not yet been defined. The rostral limit was set as the lower part of the cerebellum, the caudal limit was set as the cranial limit of C1, the dorsal limit was set as the point of intersection between the occipital bone and the cerebellum, and the ventral limit was set as the dorsal surface of the medulla oblongata. Based on these criteria, the volume of the cisterna magna was measured. In rat CLM models with cerebellar herniation through the foramen magnum, the volume of the desired CSF space, the cisterna magna, was calculated by subtracting the volume occupied by the herniated cerebellum from the total cisterna magna volume [7].

In the present study, the primary comparison showed that the FO group had a significantly greater cisterna magna volume than that of the CR group, suggesting a more pronounced effect on cisterna magna restoration with FMD alone than with FMD with cranioplasty. In 2019, Pijpker et al. [21] suggested that cranioplasty protects the enlarged cisterna magna from nuchal musculature adhesions, pressure, and extradural scarring. Contrary to these recommendations, our findings revealed that cranioplasty may impede the volume expansion of the cisterna magna achieved through FMD, challenging the previously assumed benefits of this procedure.

A recent study conducted by Ribeiro et al. [7] demonstrated that the cisterna magna volume was smaller in patients with Chiari compared to healthy individuals due to cerebellar tonsillar herniation through the foramen magnum. Additionally, the importance of the cisterna magna was emphasized, along with the necessity to compare and analyze changes in cisterna magna volume before and after surgery in patients with Chiari. However, to the best of our knowledge, no previous study has measured changes in cisterna magna volume before and after surgery in both human patients and animals with Chiari. In the current study, by utilizing a rat model of Chiari and comparing the cisterna magna volume before and after surgery, the efficacy of FMD and FMD with cranioplasty was evaluated. Therefore, this study emphasizes the rarity and novelty of measuring surgery-related changes in cisterna magna volume.

In human medicine, the extent of the craniectomy opening for posterior fossa decompression varies among surgeons, with the consensus being that sufficient occipital bone must be removed to expose the cerebellar tonsils [22]. However, an excessively wide craniectomy may lead to cerebellar herniation into the spinal canal and compression of the brainstem [23]. Cranioplasty, which repairs the bony defect caused by craniectomy, allows for broad decompression, enlarging the area around the foramen magnum and the entire posterior cranial fossa [24], which may also aid in reestablishing the proper positional relationship among the brainstem, cerebellum, and occipital bone [24]. However, the approach to craniectomy for FMD differs between humans and dogs. Dogs do not have the elongated cerebellar tonsils observed in the CM1 of humans [25]. Additionally, while the human craniocervical junction is vertically oriented, necessitating caution to prevent hindbrain migration into the craniectomy site, dogs have a horizontal orientation, reducing the likelihood of cerebellar herniation [25]. These considerations further suggest that cranioplasty, adapted from human medical practices to veterinary medicine, may not be effective in dogs.

Cranioplasty should be carefully considered; however, currently, there is no standardized procedure for cranioplasty to reconstruct the posterior cranial fossa [21]. The lack of consensus on the optimal size for craniectomy means that bone decompression and implant sizing for cranioplasty are based on non-empirical judgment and lack quantitative clinical evidence [21]. An improper design of implant castings, such as PMMA, can result in unintended placement [21]. Further disadvantages include longer surgical times, lower cost-effectiveness, and the potential for persistent untreated syringomyelia [21]. In the present study, we encountered similar drawbacks in performing cranioplasty with FMD in a rat CLM model. Due to the lack of standardized procedures for cranioplasty, there were instances when non-empirical decisions were made. Furthermore, we encountered issues regarding implant design and establishment, highlighting the need for more sophisticated techniques. Moreover, the time-consuming nature of surgery and the potential increase in postoperative complications with prolonged surgery further underscore the potential limitations associated with cranioplasty.

In recent developments in human medicine, posterior fossa decompression with duraplasty has become the most employed approach [22]. This technique involves removing sufficient occipital bone to expose the cerebellar tonsils, performing a laminectomy of the atlas, conducting an intradural inspection to identify occlusive membranes, manipulating the tonsils with or without reduction, and completing the procedure with duraplasty [22]. Similarly, our study suggested that duraplasty, alongside FMD, should be considered in veterinary and human medicine due to the beneficial outcomes thereof. Duraplasty optimally expands the cisterna magna to preserve the integrity of the subarachnoid space and ensure unobstructed CSF flow [22,26]. Moreover, it allows for the inspection and lysis of scarring or arachnoid veins at the fourth ventricle outlet, removing obstructions to CSF flow [6]. This approach is particularly effective when the arachnoid and dura are injured or enlarged, enhancing bone decompression and maximizing posterior fossa expansion [1,27]. Duraplasty can also facilitate rapid syrinx volume reduction if necessary [22,27].

This study has some limitations that should be acknowledged. This study was conducted on rats, while the primary interest in treating CLM lies in dogs, a distinctly different species. Consequently, the potential to generalize the results is limited due to physiological and anatomical variations between species. Specifically, discrepancies in cerebellar and cranial structures between dogs and rats may influence surgical outcomes and physiological responses. Therefore, caution is advised when extrapolating these experimental results to other species. The rat model may not fully represent all variables in a natural setting, necessitating consideration of the limitations of the experimental environment. Consequently, while this study offers insights into the rat CLM model, additional research is required to apply these findings to canine diseases. Generalizing the results to dogs warrants careful consideration due to interspecies differences.

## Conclusion

This study compared FMD with and without cranioplasty in a rat CLM model, providing valuable insights into surgical treatments for CLM in veterinary medicine. This study demonstrated that FMD alone and FMD with cranioplasty led to an increase in cisterna magna volume, suggesting that both methods are effective to treat CLM. Notably, FMD alone proved more effective than commonly performed FMD with cranioplasty. The superiority of FMD alone underscores the significance of directly addressing CLM structural abnormalities, facilitating CSF flow restoration and cisterna magna dilation by decompressing the foramen magnum and reducing craniospinal junction hyperdensity. Although FMD with cranioplasty aims to protect the cisterna magna from nuchal musculature pressure and extradural scarring, this study suggested that it does not offer additional benefits in preserving the cisterna magna volume. Therefore, we recommend FMD with duraplasty as an alternative approach in veterinary medicine to treat patients with CLM, potentially enhancing cisterna magna expansion by addressing and removing CSF flow obstructions. Future research should focus on unraveling the underlying mechanisms of CLM and refining surgical techniques to improve patient outcomes.

## Acknowledgments

The Veterinary Medical Teaching Hospital of Konkuk University, Republic of Korea, supported this study.

## Supporting information

**S1 Fig. Comparison of the cisterna magna volume among the groups.** Post hoc analysis indicated that cisterna magnum volumes were significantly greater in the FO (foramen magnum decompression [FMD] only) group than in the control group (p < 0.001), in the CR (FMD with cranioplasty) group than in the control group (p = 0.003), and in the FO group than in the CR group (p = 0.005).

S1 Information. A concise summary of Chiari-like malformation in dogs S2 Information. Structural features of the cisterna magna in humans

